# Application of a Novel Numerical Simulation to Biochemical Reaction systems

**DOI:** 10.1101/2023.08.10.552732

**Authors:** Takashi Sato

## Abstract

**Motivation:** Omics data and single-cell analyses have recently produced many biological informatics. These require simple, fast, and flexible numerical/analytical methods such as ordinary differential equations. However, formulating these equations and their computational processes `can` be expensive and imprecise for simulating reactions involving genes and a small number of molecular systems. Therefore, developing a straightforward simulation method is necessary.

**Results:** We developed a natural number simulation (NNS) method using binomial probability-based stochastic algorithms. Hence, this paper simulated one-gene systems for feedback and feed-forward reactions, allosteric biochemical reactions, and SIR-type population dynamics. Furthermore, NNS can calculate any biological reaction systems written using stoichiometric formula. Thus, NNS provides a comfortable simulation tool for the scientific and engineering fields; algorithms and applications are detailed using Python.

**Availability and implementation:** Calculation results and the program are available as supplementary information in binomial_v15.zip in https://binomial-simulation.com/en/python-program/.

**Supplementary Information:** Supplementary data are available in this pdf file.

**Graphical Abstract:** 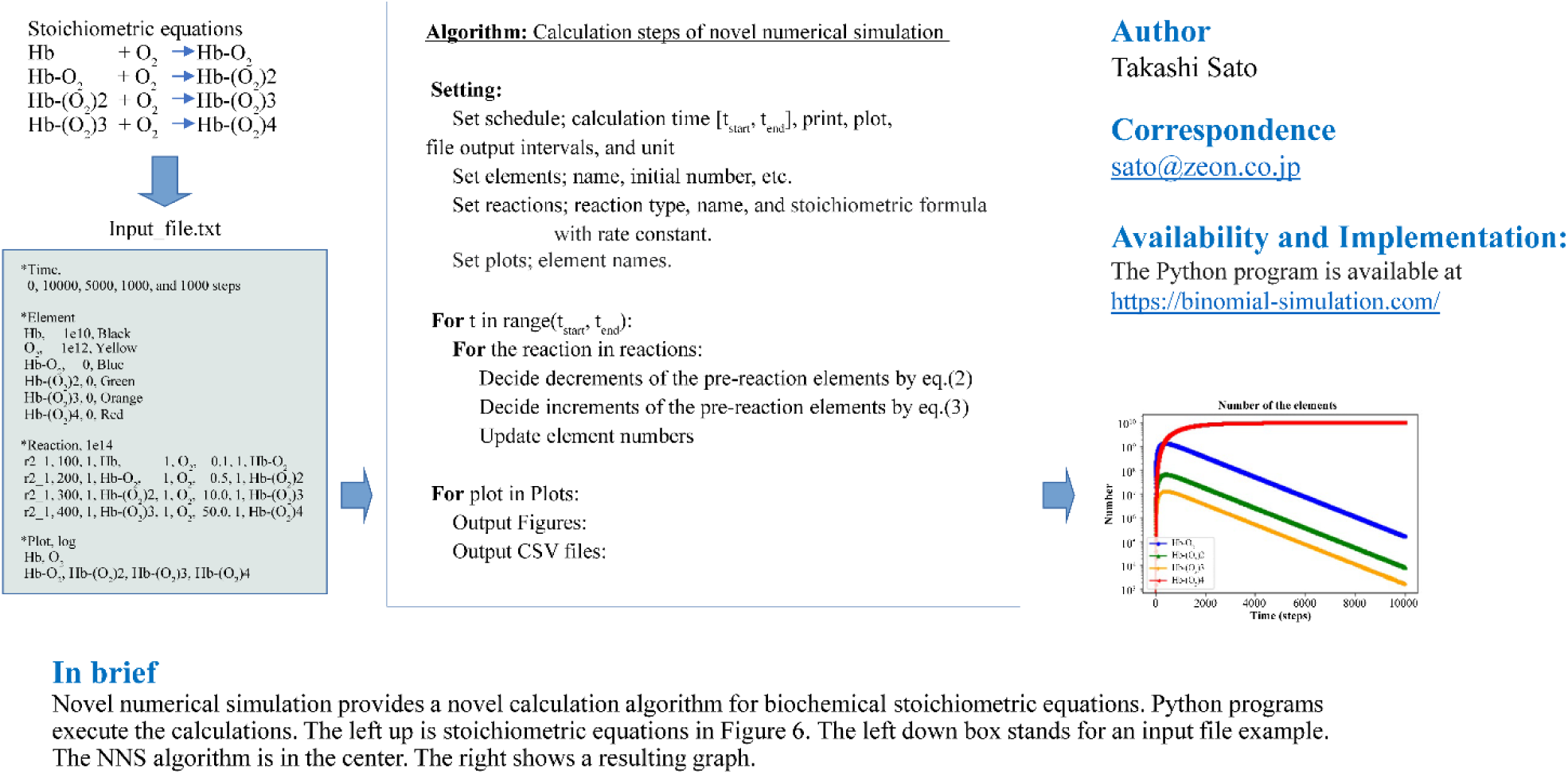

## Introduction

This paper presents a simulation program for biological reaction systems using the natural number simulation (NNS) method. NNS models biochemical reactions in a stoichiometric manner and calculates them on time series. It provides a simple modeling technique and fast calculations. Various simulation and analysis tools have been developed for modeling biological systems, incorporating results from other fields (Huizing *et al*. 2022, Browning *et al*. 2022, Blinov *et al*. 2004). However, they may not be suitable for complex biological systems due to the costs of formulating equations and numerical calculations (Shaikh *et al*. 2022, Brinkrolf *et al*. 2021). Therefore, user-friendly tools are needed.

NNS, implemented in Python, offers a convenient package for simulating and analyzing complex biological systems. It uses the probabilistic binomial distribution to calculate reactions on natural numbers. The calculations occur in one decreasing and increasing process per reaction. While ODEs and Gillespie’s algorithm have been widely used for simulating reaction systems, they may not be optimal for complex biological systems (Alon 2019, Gillespie 1977). Alternatively, NNS accurately represents the small numbers of molecules involved in DNA-related reactions and enables numerical simulation.

Transcription and translation from DNA are examples of NNS application (Watson *et al*. 2013). These reactions proceed specifically and selectively, and NNS can accurately describe them using stoichiometric equations with a rate constant. By providing solutions in terms of natural numbers, NNS eliminates uncertainty in DNA, mRNA, and protein quantities.

Furthermore, the binomial distribution of the NNS algorithm allows for the easy calculation of information such as Shannon’s entropy (Adriaans and Benthem 2008, Shannon 1948). A calculation sample is provided in the *Discussion*.

## System and methods

### Overview

The proposed approach provides time-developing solutions for calculating the stoichiometric reaction using a binomial algorithm. The general stoichiometric formula is as follows:

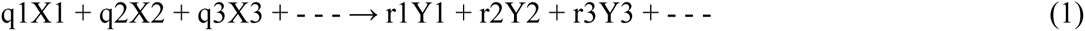

with rate constant k, X1, X2, ---denote molecular elements before the reaction, while Y1, Y2, ---are molecular elements after the reaction. The parameters q1, q2, ---, r1, r2, ---are reaction’s orders. The rate constant k determines the probability of creating new materials Y1, Y2, ---. The reaction orders are also natural numbers if the elements contain a natural number of atoms in each molecule. This approach deals with stoichiometric reactions but does not restrict the precise formula of chemical reactions. This means we can describe Equation (1) as a model formula with the conservation of atom numbers.

After formulating Equation (1), the initial numbers for each element are needed to calculate the time sequence of the element’s numbers. For example, suppose Xi_n is defined as the elemental number of Xi, and qi is the order; then, the following calculation provides the decrement and increment quantities of the elements.

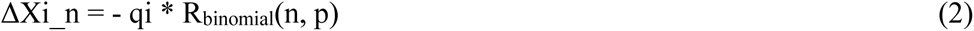

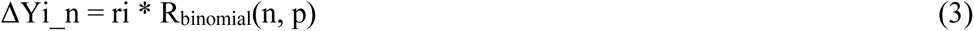

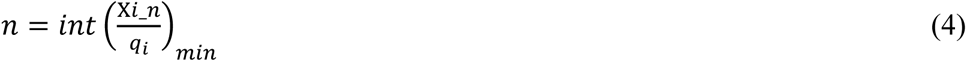

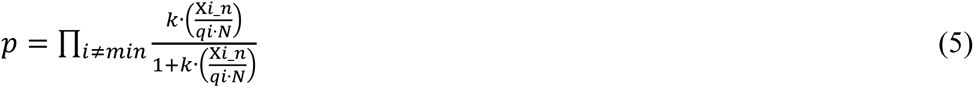

R_binomial_ (n, p) is a random-number-generating function that returns a natural number using binomial distribution: *n* is a trial number with binary success and fault values, and *p* is the probability of success. NumPy’s random.binomial(n, p) function is used in this program. The function returns one successful natural number under *n*, including zero. Stochastic simulation often uses these discrete numbers (Székely Jr. *et al*. 2014, Gholami *et al*. 2021).

The int(Xi_n/qi)_min_ denotes the smallest number among the values int(Xi_n/qi) of the previously introduced elements. Note that int(Xi_n/qi) is a rounded down to the nearest integer of Xi_n/qi, the reaction unit in the reaction formula. Further, *N* in Equation (5) is a normalization parameter that determines the system size in a concerned reaction system. At the same time, *p* is the mathematical product of each function with the number of elements Xi_n, order qi, rate constant *k*, and normalization parameter *N*. Equation (5) does not contain the selected element in Equation (4). The constant’s units differ from each other in ODEs because of the definitions of the terms (Soliman *et al*. 2010, Mangan *et al*. 2016). Conversely, the value *k* in NNS is nondimensional because NNS formulates only numbers of molecules and/or elements. This would facilitate relative comparisons for each reaction.

Equations (4) and (5) have some minor formulas. One is in the case of a one-element reaction, as follows:

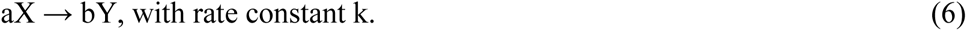

X and Y are molecular elements, and *a* and *b* are orders. The values of *n* and *p* are derived from the following formulas

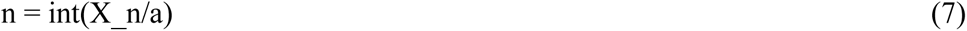

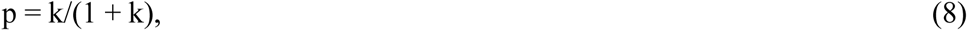

where X_n is the number of the element X. Equation (7) is similar to Equation (4). However, *p* in Equation (8) has a simpler formula because Equation (6) has only one type of element before the reaction.

Another exceptional reaction type is as follows:

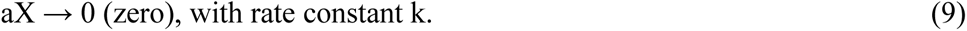

This expression indicates that element X is decomposed and disappears, so Equation (3) does not work. Thus, only X decreases with Equations (7) and (8).

Another type of reaction defines linear decreasing and increasing reactions for one element. One needs these reactions for the mathematical formulation of linear changes via molecular addition and subtraction. The linear decreasing case is defined similarly to Equation (9):

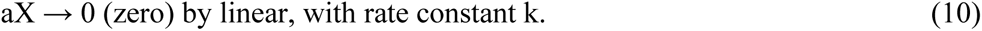

Moreover, the increment of X is determined by the following formula:

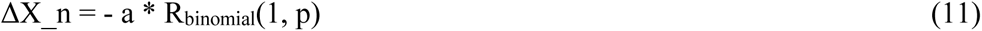

where *p* is the same as in Equation (8), the definition of n = 1 reads as linear decreasing. Subsequently, the linear increasing case is defined by the next reaction formula

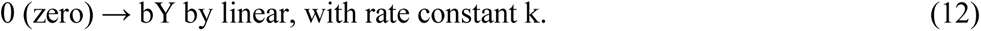

The next formula also determines the increment of Y

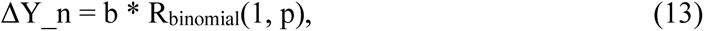

where *p* is the same as in Equation (8).

A biological system has many incredible reactions (Nelson *et al*. 2021). Increments of elements by those reactions can be described by (1), (6), (9), (10), and (12), with each reaction being easily defined if the before and after elements, their orders, and rate constants are known.

An input file is executed in Spyder by writing the file-name line in the main program file (i.e., binomial_v015.py), fName = “inp_file.txt.” The “Run” command in Spyder executes the calculation. After running, results files are created in the folder “inp_file.txt.” Another execution is done in the command line. For example,

$ python binomial_v015.py inp_file.txt

This line executes the calculation, giving similar results as above. Notably, the input file is in the same program files folder or another one level up of the program files. One text input file needs to define calculation time, elements, reactions, and plots (Table 1).

**Table 1.**
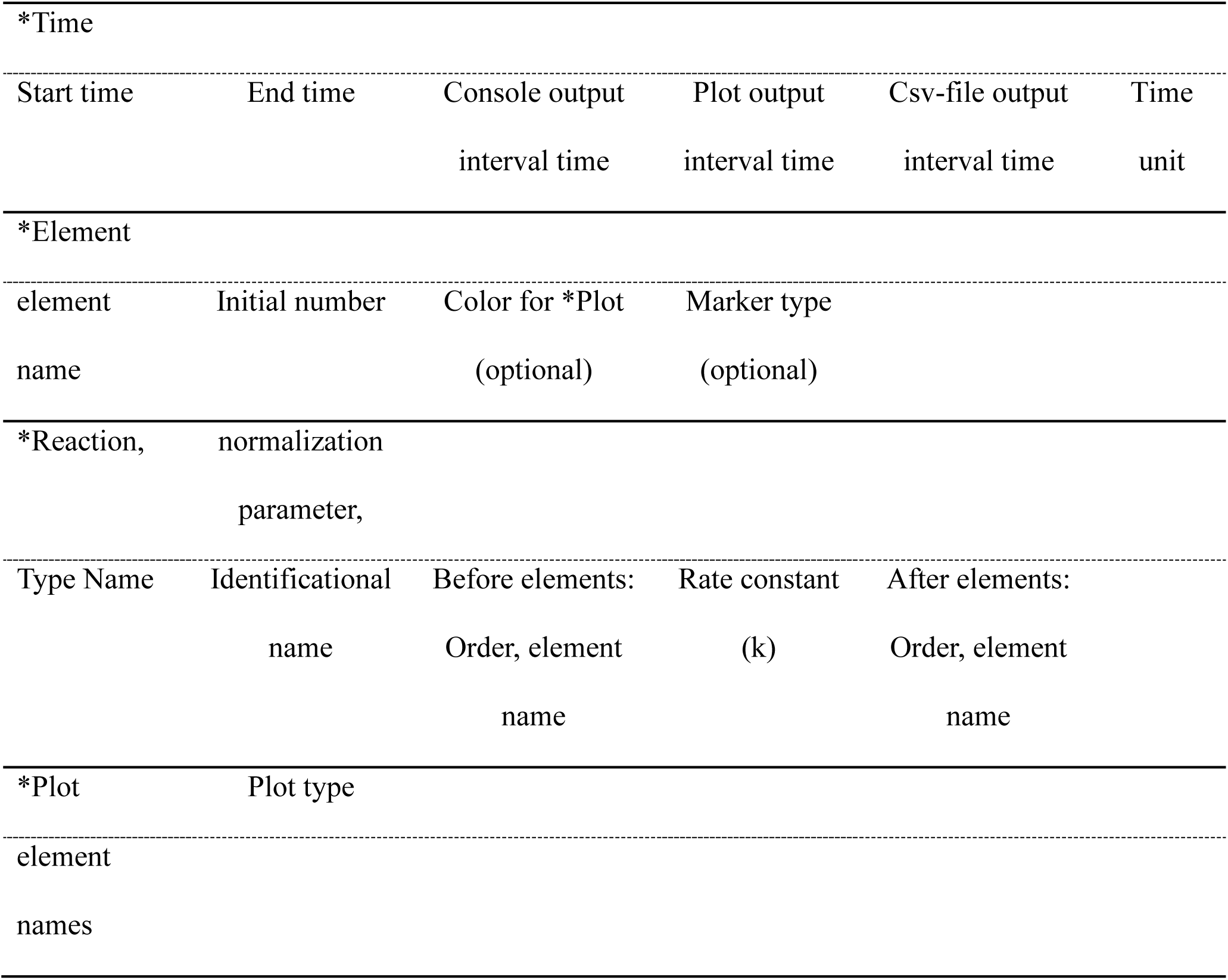
Standard input file style.

Additionally, since the reaction statement, *Reaction is important, many definitions were prepared (Supplementary Tables S1 and S2). Those definitions included the above formulas to calculate reaction increments.

We conducted biological systems’ experiments with the in-flow and out-flow of metabolic molecules, signal proteins, etc. These element definitions are made available (Supplementary Table S3), providing a wide range of simulations for biological reactions. A Python program made by the author is available (Sato, 2022).

### Algorithm and results

The calculation schematic flow and detailed algorithms for element increments are depicted in Figures 1 and Supplementary Figure S1, respectively. The simplicity of the procedure is demonstrated through six examples using corresponding input files.

**Figure 1.**
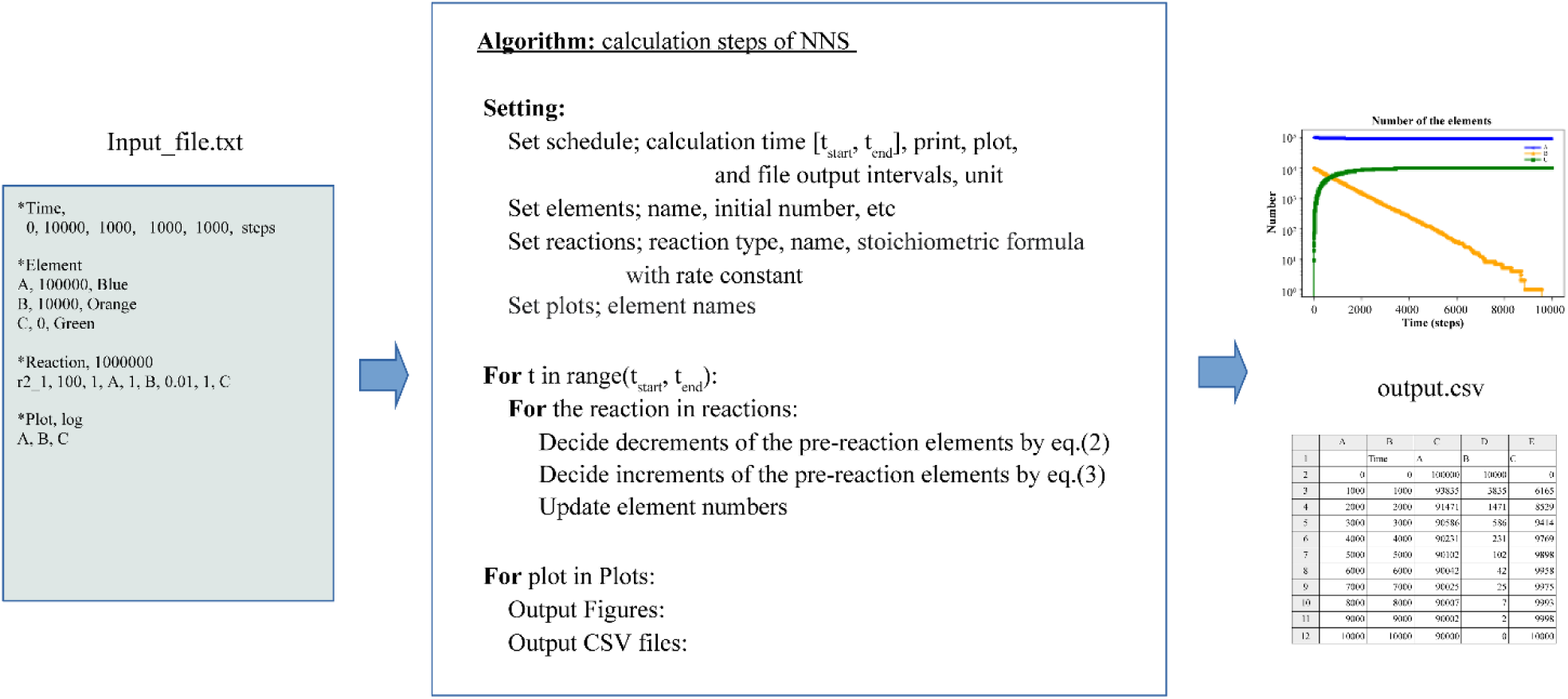
Calculation flow and algorithm. (Left) Input file example. (Center) NNS algorithm. (Right) Graph and CSV file results.

### Simple reaction model

Consider a simple reaction:

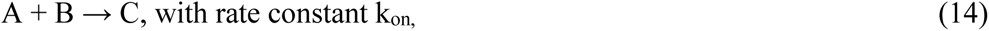

where A and B may be two protein molecules that irreversibly bind and become molecule C, the rate constant k_on_ determines the reaction speed per time.

Figure 2a shows an input file, which contains four items, *Time, *Element, *Reaction, and *Plot, as shown in Table 1. One reaction, r2_1_100 (“r2_1, 100” in the input file) is defined. Calculation time starts from zero and ends at 10000. The algorithm proceeds step-by-step by one-time unit. The unit of time is nondimensional in calculations. So, the unit of time should be determined by the user according to the characteristics of the user’s calculation target. NNS uses natural numbers so that even though the number of element B and C is small, the solution will give the exact number of elements (i.e., one, two, three, etc.), as plotted in Figure 2b.

**Figure 2.**
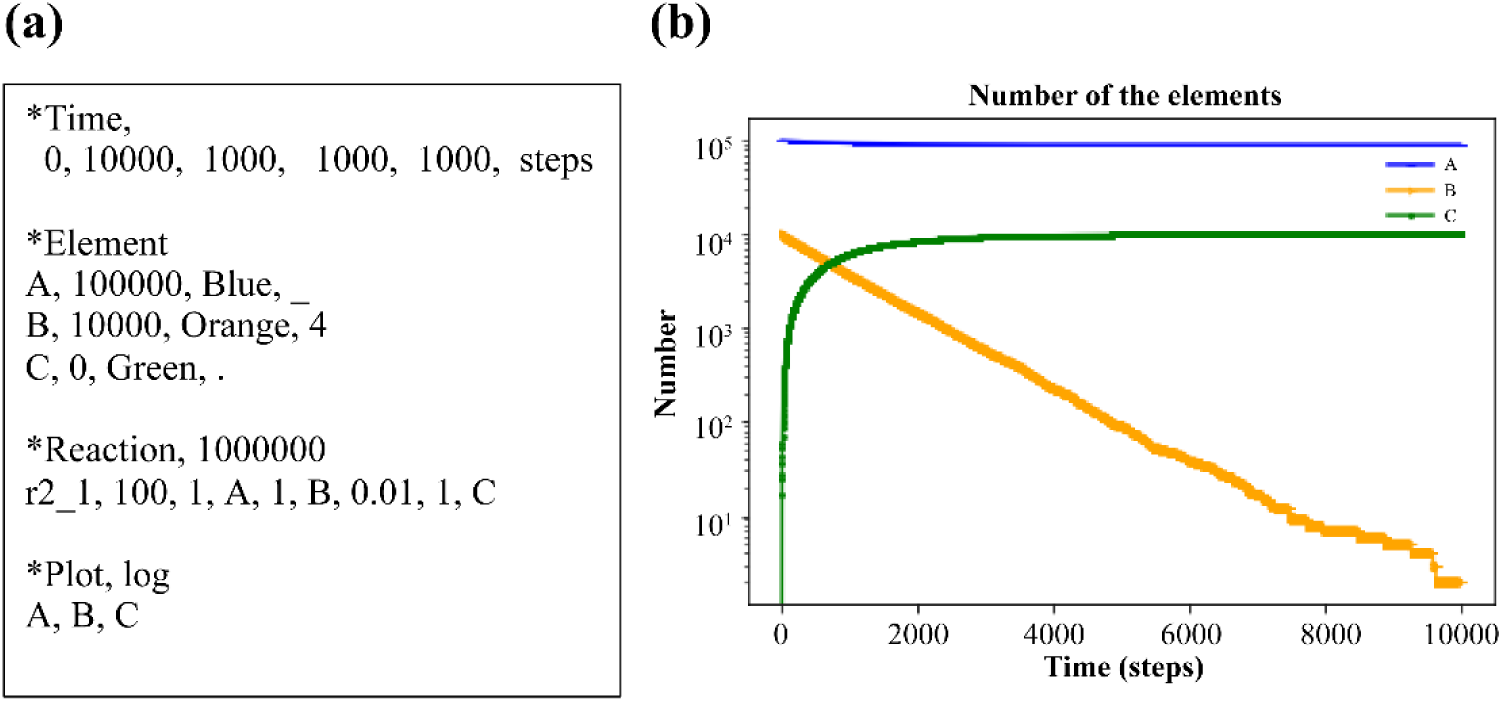
(a) Input text and (b) calculation result.

In contrast, Figure 3a shows another input file with inverse reaction r1_2_200.

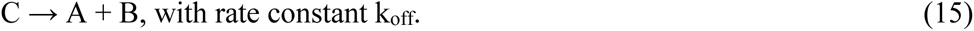

**Figure 3.**
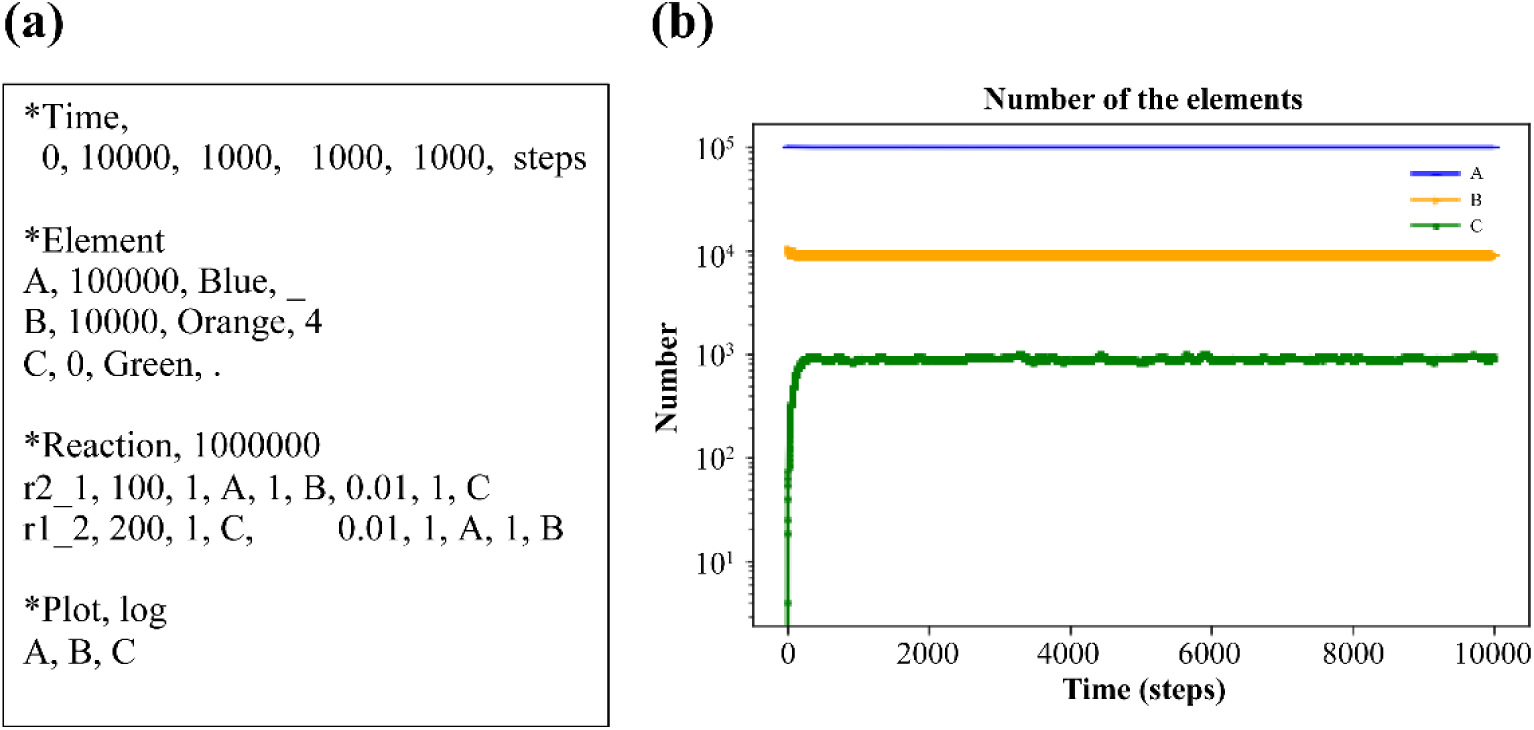
(a) Input text. (b) Calculation result.

Biological systems have many such reactions (Craig *et al*. 2021). The rate constant k_off_ differs from k_on_ in Equation (14) due to the thermodynamic principle in the system. Figure 3b shows the equilibrium numbers of A, B, and C.

### Michaelis–Menten model

Michaelis–Menten kinetics is applied using NNS (Gilbert *et al*. 2006). In Figure 4a, E is an enzyme, S is the substrate, ES is the enzyme–substrate complex, and P is the product (Craig *et al*. 2021). The corresponding scheme is as follows:

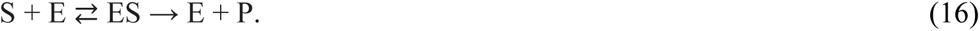

**Figure 4.**
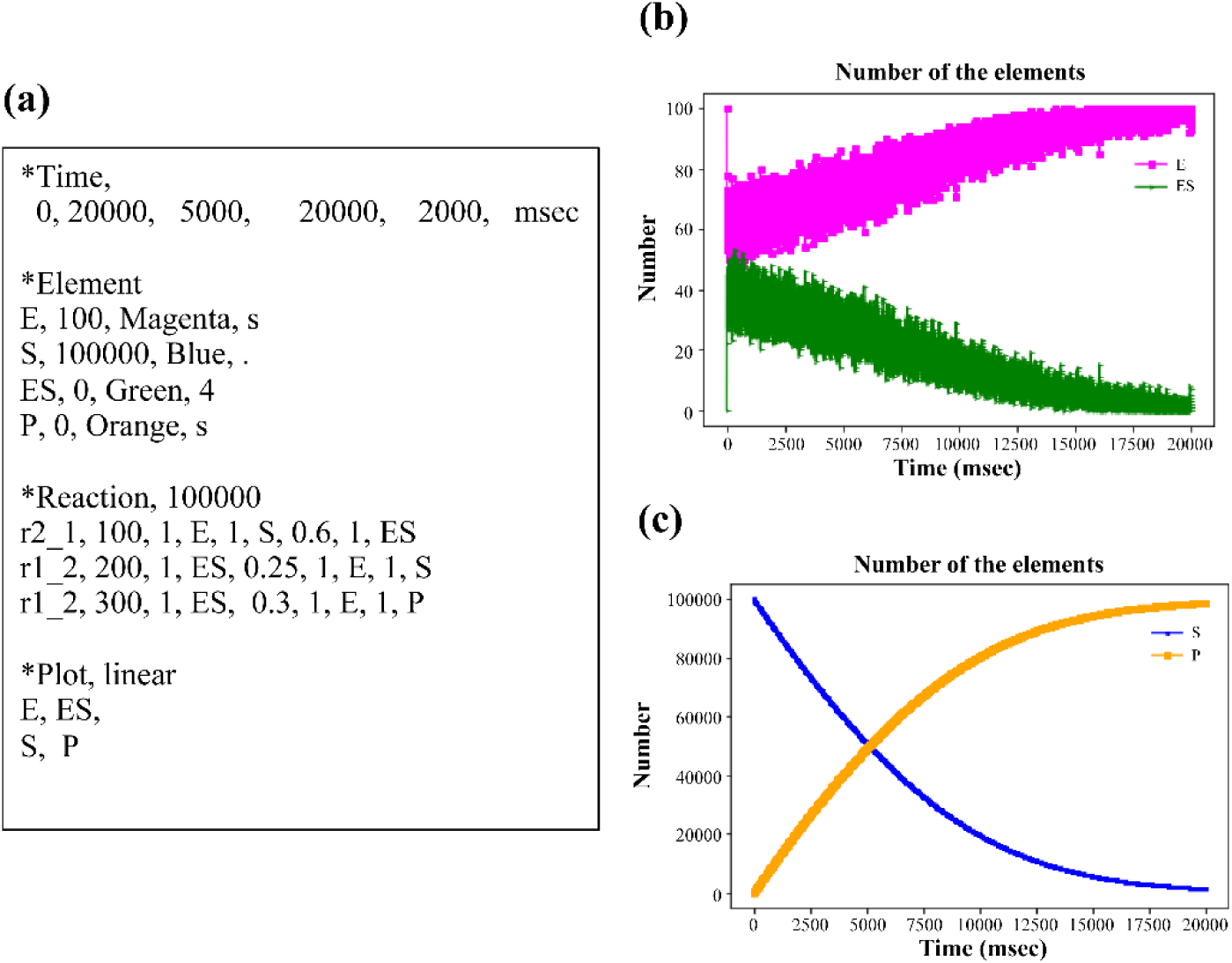
Michaelis–Menten kinetics. (a) Input text. (b) Results of E and ES. (c) Results of S and P.

NNS defined three reactions in the input file. The total E number (E_n + ES_n), is constant though E and ES fluctuate in time series (Figure 4b). Furthermore, S decreases with increasing P (Figure 4c).

### Monod–Wyman–Changeux allosteric model

NNS can handle allosteric models (Goodey and Benkovic 2008, Machado *et al*. 2015, Wodak *et al*. 2019). The allosteric transition of hemoglobin has one of the most interesting behaviors because of molecular adaptation in vertebrates (Eaton 2022, Hiroshi 2008). Supplementary Figure S2a presents an input file for oxygen binding to hemoglobin at different rate constants (0.1, 0.5, 10, and 50). However, Supplementary Figure S2b contains same four rate constants (0.1, 0.1, 0.1, and 0.1). Figure 5a shows the results with the allosteric effect, the increasing rate of Hb-(O_2_)4 was higher than from the no allosteric effect model of Figure 5b at early time (Supplementary Figure S3a). Moreover, the ratios of Hb-(O_2_)4 to other complexes, Hb-(O_2_), Hb-(O_2_)2, and Hb-(O_2_)3 became larger than from the no allosteric effect model of Figure 5b at any time (Supplementary Figure S3b)).

**Figure 5.**
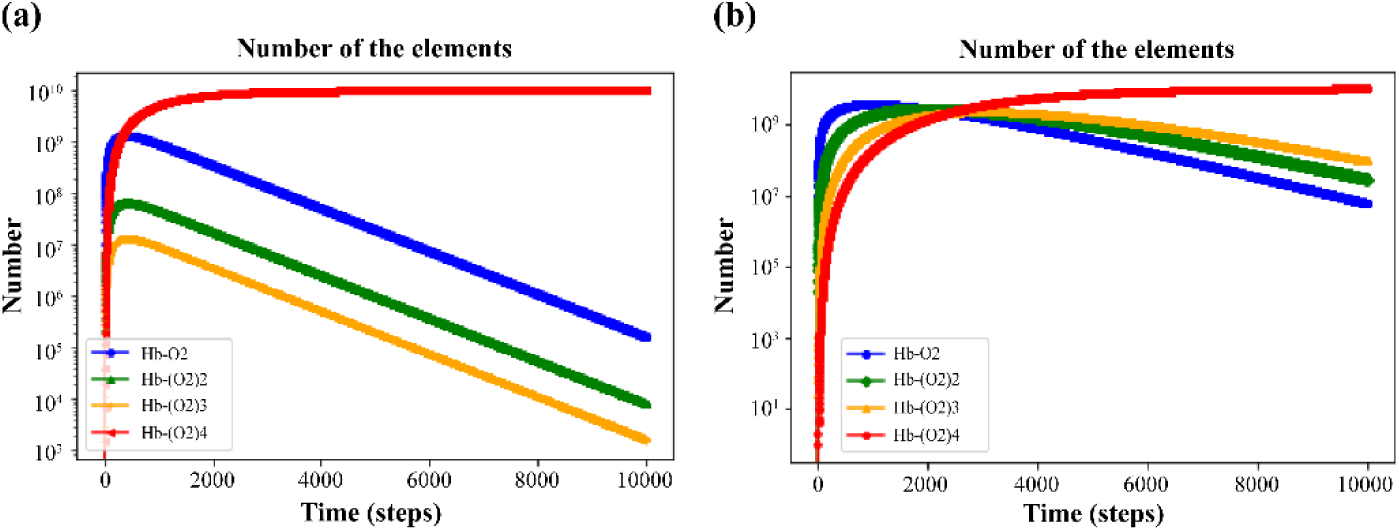
Results of Monod–Wyman–Changeux allosteric model. (a) Result of Supplementary Figure S2a, with rate constants; 0.1, 0.5, 10, and 50. (b) Result of Supplementary Figure S2b, with rate constants; 0.1, 0.1, 0.1, and 0.1.

### Feedback loop model

NNS supports the modeling of feedback loops in biological systems. Feedback loops often occur to regulate biological systems (Alon 2019, Ferrazzi *et al*. 2011). Figure 6 shows a model in which stimulus material activates DNA. DNA is transcribed into mRNA, which is then translated into protein. The protein binds to the DNA to deactivate and turn it into DNA_d (deactivate). Stoichiometric formulas are available on some modeling hypotheses. Supplementary Figure S4 shows an input file, which models that ribonucleotide is one type and amino acid is also one type. It is advantageous that NNS can describe one DNA molecule and can accurately deliver the corresponding property if a precise description is required.

**Figure 6.**
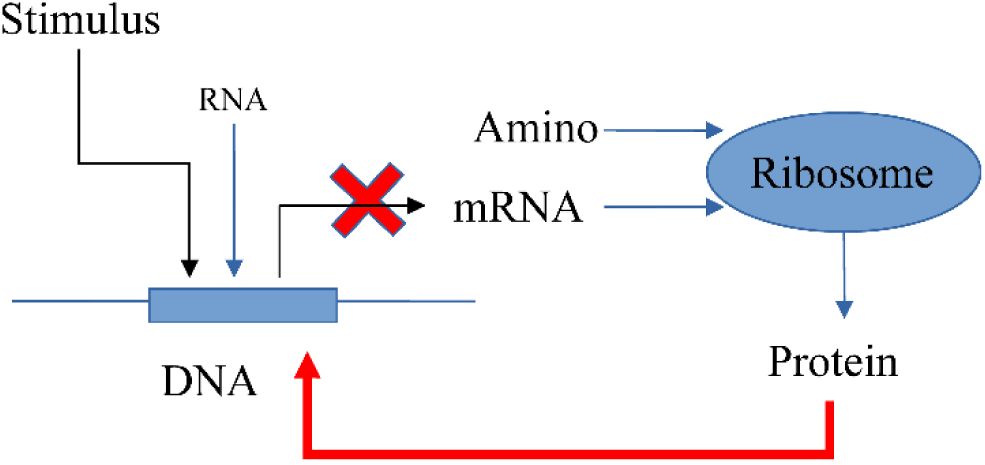
Feedback system model for one gene DNA. Stimulus: protein molecules bind DNA; DNA: one gene; RNA: some ribonucleic acids; mRNA: messenger-RNA; amino: some amino acids; ribosome: ribosome for translation; and protein: a protein made by ribosome.

Figure 7 shows the result with feedback, while Supplementary Figure S5 shows the result without feedback, where the reaction r2_1_500 is deactivated. The number of produced proteins is kept low in systems with feedback control because of the protein-deactivated DNA.

**Figure 7.**
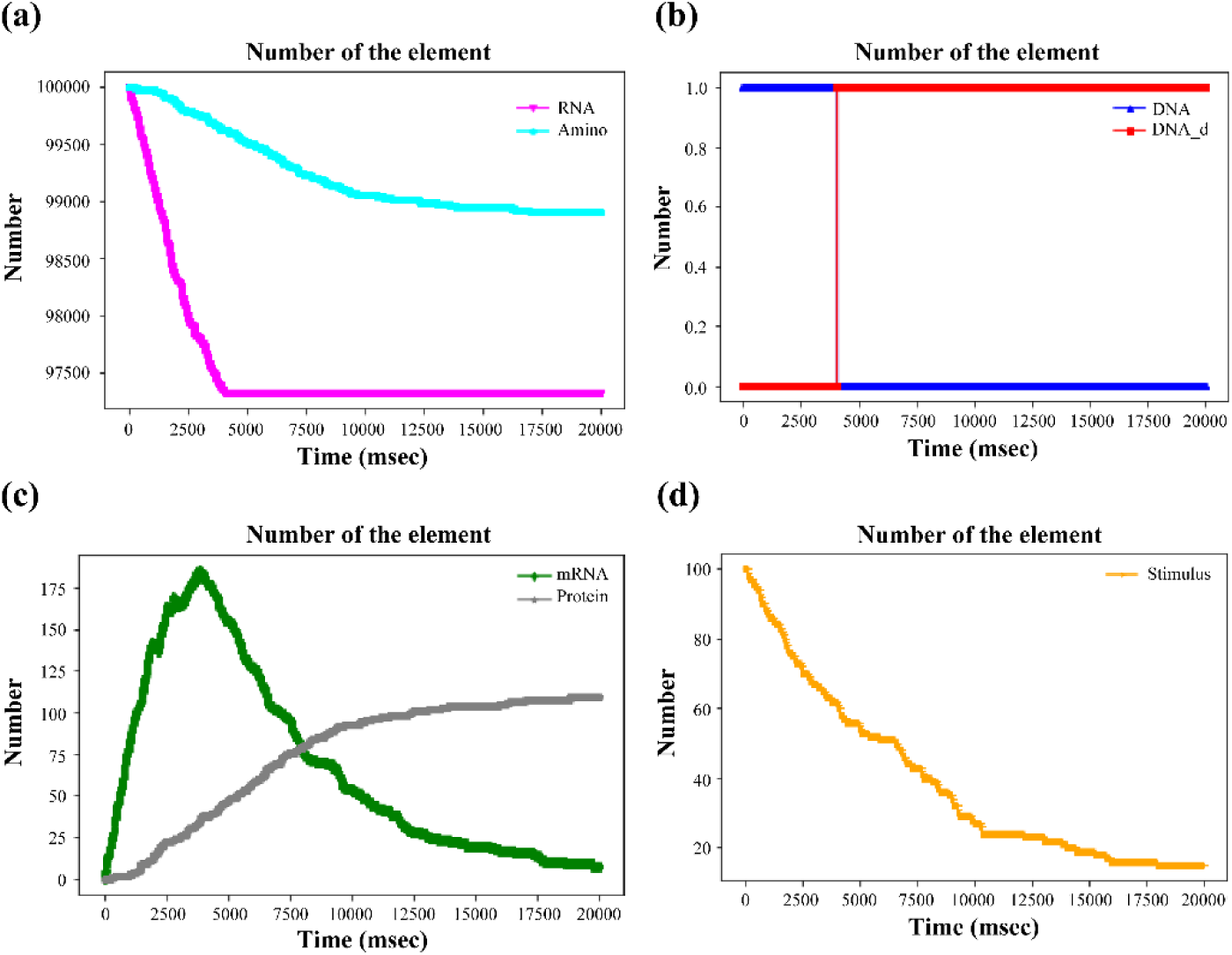
Results of the feedback system of the input file in Supplementary Figure S4. (a) RNA and amino acids. (b) DNA and DNA_d. (c) mRNA and protein. (d) Stimulus.

### Feed-forward loop in a biological system

The feed-forward loop is another common biological system (Duc-Hau Le *et al*. 2011, Kremling *et al*. 2008, Macía *et al*. 2009). Figure 8 depicts a schematic reaction process (Alon 2019). Stimuli Sx and Sy ultimately promote protein pZ production. Many elements and reactions are contained in the system; pX, pY, and pZ are proteins, pX* and pY* are activated types of each protein, “amino” is an amino acid, DNA-Y, and DNA-Z are DNA molecules corresponding to proteins pY and pZ, respectively. Furthermore, DNA-Z_X*, DNA-Z_X*_Y*, and DNA-Y_X* are activated DNA molecules, i.e., DNA-Z_X* stands for activated DNA-Z by activated protein pX*, DNA-Z_X*_Y* for activated DNA-Z by activated proteins pX* and pY*, and DNA-Y_X* for activated DNA-Y by activated protein pX*.

**Figure 8.**
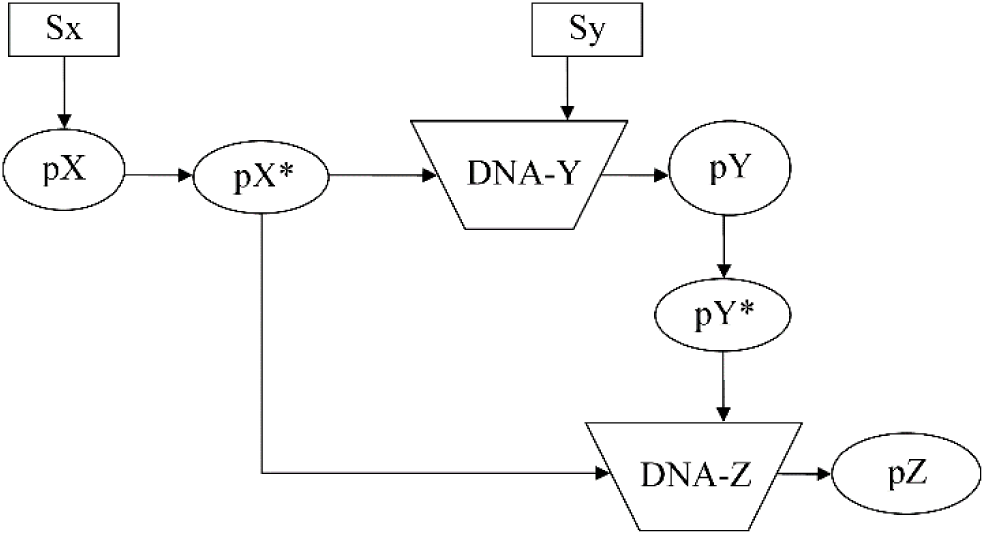
Feed-forward system. Sx and Sy: Stimuli; pX, pY, and pZ: proteins; pX* and pY*: activated proteins of pX and pY; DNA-Y and DNA-Z: DNA for each protein pY and pZ.

Supplementary Figure S6 presents the reaction definitions. They contain four types of reactions, r2_1, r2_3, and r3_3 (shown in Supplementary Table S1) and rM (shown in Supplementary Table S2). Figure 9 shows the results of this system, i.e., pZ increases after pY, and pY* increases after time 4000.

**Figure 9.**
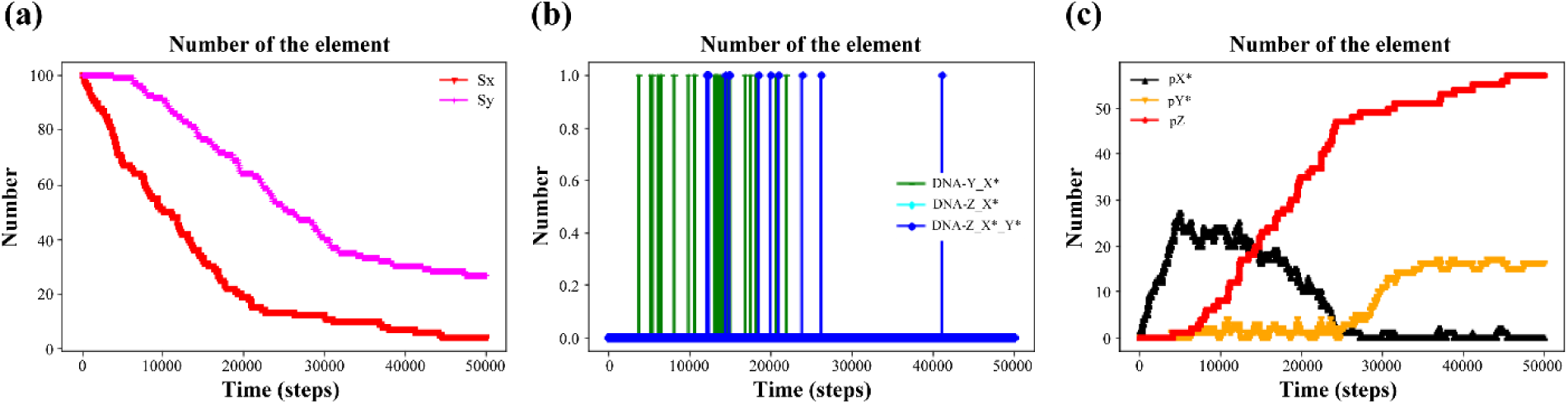
Results of the feed-forward system in Supplementary Figure S6 until time = 50000. (a) Sx and Sy. (b) DNA-Y_X*, DNA-Z_X*, and DNA-Z_X*_Y*. (c) pX*, pY*, and pZ.

Supplementary Figure S7 changed the Sx and Sy definitions, i.e., Sx is added at time 5000 and Sy at 10000 using *ElementInOut. Supplementary Figure S8 shows the corresponding results, i.e., pX* and pY* shifted, then pZ appeared around 10000. Some elements may be added at any time in an *in vivo or in vitro system*. It is easy to describe such situations with NNS, as it can perform modeling flexibly. Hence, analyzing experimental data using NNS could provide an appropriate interpretation of these at the gene level.

### SIRS and SEIRS model

The SIR model (Murray 2002) is a basic model for infection dynamics, with S denoting “susceptible,” I denote “infected,” and R denoting “recovered.” There are many types of variations. For example, the SEIRS model considers “exposed” E and simulates recruitment into S (Bjørnstad 2020). Supplementary Figure S9 shows three models: SIRS, SEIRS, and SEIRS with birth and death. In model (c), S1_sum stands for the summation of birth, and I_d (death) stands for the summation of infected patients that died. Supplementary Figure S10 shows the corresponding results, illustrating the simulation’s extensibility to different scenarios.

## Discussion

The properties of a stochastic process with probability parameters can introduce information and entropy. Entropy is a thermodynamic, statistical, and information theory concept (Baez and Pollard 2016). The parameters *n* and *p* in np.random.binomial(n, p) is determined from the reaction definition in the overview of the *System and methods*. Reaction conditions determine these numbers at the time. Accordingly, information on reaction can be defined as shown below:

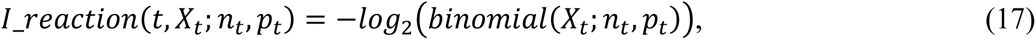

where binomial(*X*; *n*, *p*) is a binomial distribution function with a stochastic variable *X* determined using *n* and *p* of the np.random.binomial(*n, p*) at each reaction step *t* (Chanda *et al*. 2020). The *I*_*reaction* becomes a large number when the probability binomial(*X*_*t*_; *n*_*t*_, *p*_*t*_) is small. Supplementary Figure S11 (a, b, and c) illustrates the trends of information on reaction (I_reaction) with respect to the stochastic variable X. Supplementary Figure S11d shows how maximum and minimum information values change with *n* at a fixed *p*. Large information values indicate the rare occurrence of the reaction. I_reaction provides insight into the reaction dynamics, and its value continues to increase as the reaction progresses.

When the variables *p* and *n* are fixed, the binomial(X; n, p) distribution generates values for the stochastic variable X. Then, the reaction entropy (RE), similar to the Shannon entropy, can be defined as follows (Ben-Naim 2012):

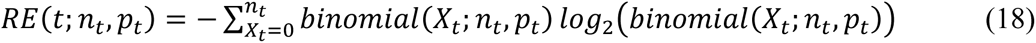

where summation Σ indicates the sum of stochastic variables *X_t_* from *X_t_ = 0* to *X_t_ = n_t_* as discrete natural numbers. RE is determined using *n_t_* and *p_t_*, as shown in Supplementary Figure S12. Equation (18) describes a state quantity, which is similar to a thermodynamical variable, depending on *n_t_* and *p_t_* at the time.

For example, considering a DNA-type reaction and the feedback model in Supplementary Figure S4, variable *X_t_* is chosen randomly per reaction at that time as the reaction proceeds. Once the number is decided, the information of Equation (17) can be calculated. Here, the accumulated information on reaction until time *t* is defined as follows:

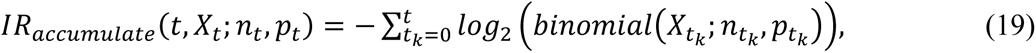

The summation means the sum of the information on reaction from time 0 to time *t*. The *IR*_*accumulate*_(*t*, *X*_*t*_; *n*_*t*_, *p*_*t*_) implies generated information of each reaction until the time reaches the value *t* in the system. Supplementary Figure S13 shows the results of Equation (19) in the case of a feedback loop of Supplementary Figure S4 comprising five reactions. The IRaccumulate values reflect the importance and diversity of biochemical reaction processes.

Conversely, RE represents the randomness and activity of reactions (Baez and Pollard 2016, Roach 2020, Uda 2020). Supplementary Figure S14 shows the REs of the five reactions in Supplementary Figure S4. RE showcases how the activity and randomness change over time, and it serves as a measure of reactivity.

Information on reaction and RE are crucial concepts for analyzing reaction properties. They provide insights into the changes and characteristics of reactions, allowing for a deeper understanding of biochemical systems.

## Conclusions

From this study, the following six conclusions are obtained:

1. NNS is a novel algorithm that uses exact numbers and provides a time-developing solver for stoichiometric reaction systems.
2. Simple binding reactions and their equilibrium were successfully calculated.
3. Important biological reactions, such as Michaelis–Menten kinetics and Monod–Wyman– Changeux allosteric model, were performed.
4. Feedback with one gene and feed-forward systems with two genes were analyzed, demonstrating their regulatory effects.
5. SIR-type models with additional characteristics were successfully simulated.
6. Information on reaction and RE was obtained using the NNS formula.

The NNS approach complements existing methods based on ordinary differential equations (ODEs) and contributes to the comprehensive study of complex biological systems.

Biological systems exhibit nested structures and intricate interactions, such as symbiosis and cell– cell interactions. For example, multicellular organism contains organs, cells, cytoplasm, and nucleus. Furthermore, they perform symbiosis between different types of life forms, for example a human and E. Coli, and cell–cell interactions, such as between antigen presenting cell and T cell inside a life. To model such complex systems using NNS, the following approaches will be pursued:

1. Gathering stoichiometric reaction data for a comprehensive database.
2. Developing a membrane system model to represent cells, organs, and organelles using the concept of “set” for separation (Supplementary Figure S15a). The “set” has a membrane system to transport substrates for biomolecules inside-out and reverse.
3. Creating a multilayer model to represent cells in a multicellular organism using the “set” concept (Supplementary Figure S15b).
4. Developing a direct-interaction model between sets (Supplementary Figure S15c).
5. Combining multilayer structures and direct interactions within sets, incorporating reactions within these sets (Supplementary Figure S15d).

These future research directions will enhance the capabilities of NNS and enable more sophisticated modeling of biological systems.

## Acknowledgments

I want to thank my colleague, Kunihiko Oishi, Chika Morimoto, Minami Sakamoto, and Tatsuro Omori in my company team.

## Data availability

The data are available at https://binomial-simulation.com/. In addition, a zip file, binomial_v015.zip, contains the results of this paper.

## Author Contribution

The author confirms sole responsibility for the following: study conception and design, data collection, analysis and interpretation of results, software and manuscript preparation.

## Supplementary Materials

### Supplementary Tables

**Table S1.** Standard reaction types and definition styles for NNS.

| Type | Identification | Before element: | Rate | After element: | Equation |
| --- | --- | --- | --- | --- | --- |
| name | name | Order, element name | constant | Order, element name |  |
| r1_0 | name | q1, X1, | k, | None | eq(9) |
| r1_1 | name | q1, X1, | k, | r1, Y1 | eq(6) |
| r1_2 | name | q1, X1, | k, | r1, Y1, r2, Y2 | eq(1) |
| r1_3 | name | q1, X1, | k, | r1, Y1, r2, Y2, r3, Y3 | eq(1) |
| r2_1 | name | q1, X1, q2, X2, | k, | r1, Y1 | eq(1) |
| r2_2 | name | q1, X1, q2, X2, | k, | r1, Y1, r2, Y2 | eq(1) |
| r2_3 | name | q1, X1,q2, X2, | k, | r1, Y1, r2, Y2, r3, Y3 | eq(1) |
| r3_1 | name | q1, X1, q2, X2, q3, X3 | k, | r1, Y1 | eq(1) |
| r3_2 | name | q1, X1, q2, X2, q3, X3 | k, | r1, Y1, r2, Y2 | eq(1) |
| r3_3 | name | q1, X1, q2, X2, q3, X3 | k, | r1, Y1, r2, Y2, r3, Y3 | eq(1) |
| r1_- | name | q1, X1 | k, | 0, 0 | eq(10) |
| r1_+ | name | 0, 0 | k, | r1, Y1 | eq(12) |

**Table S2.**
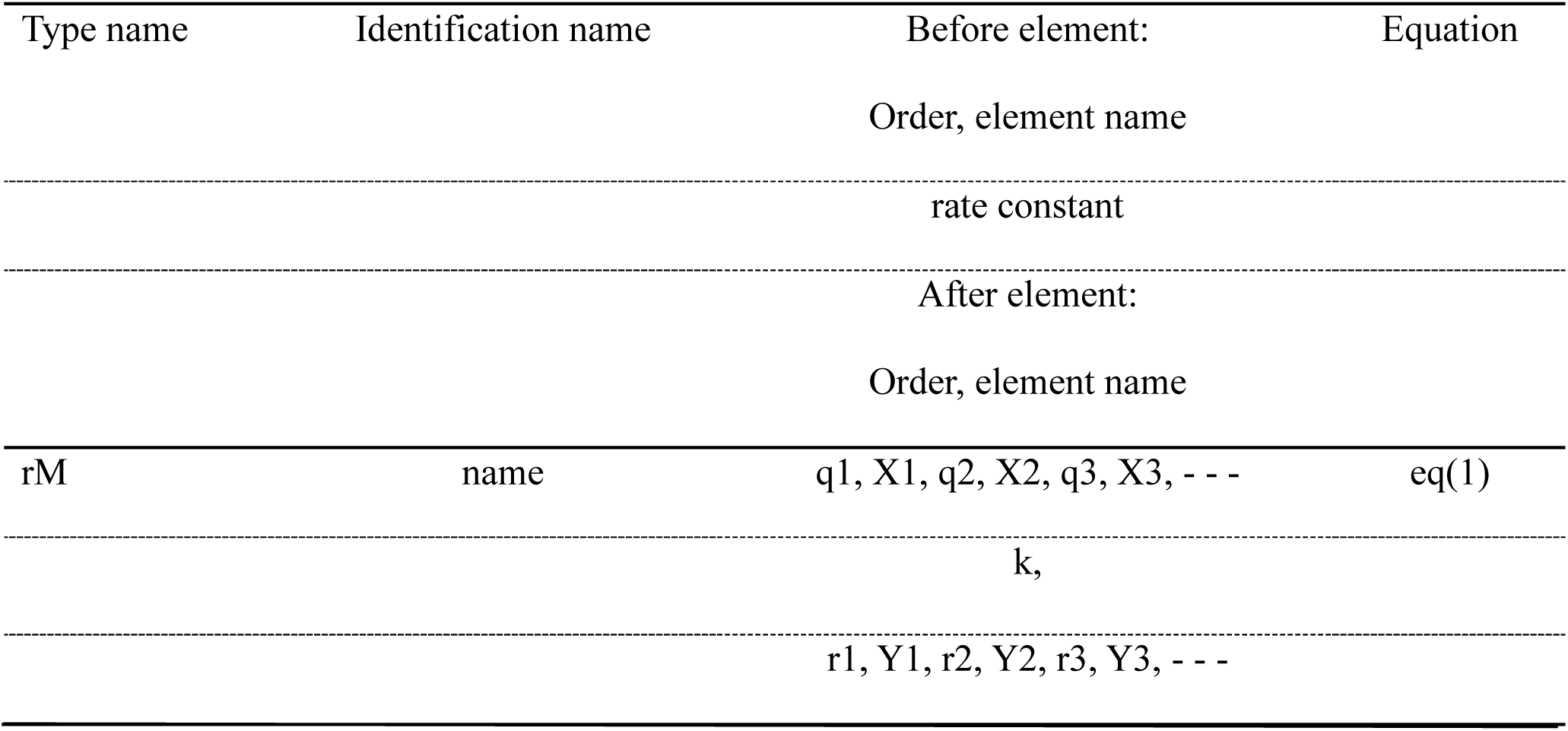
rM-reaction definition style for NNS.

**Table S3.** Additional statements on the input file, *ElementInOut.

| *ElementInOut |  |  |  |  |  |  |  |  |  |  |
| --- | --- | --- | --- | --- | --- | --- | --- | --- | --- | --- |
| Element | Initial | Color | Marker | Element | type- | A | Amount | A | Amount | And |
| name | number | for | type | type | 0 | time |  | time |  | so |
|  |  | *Plot |  | (optional) |  |  |  |  |  | on |
| Element | Initial | Color | Marker | Element | type- | Interval | Amount |  |  |  |
| name | number | for | type | type | 1 | time |  |  |  |  |
|  |  | *Plot |  | (optional) |  |  |  |  |  |  |

### Supplementary Figures

**Figure S1.**
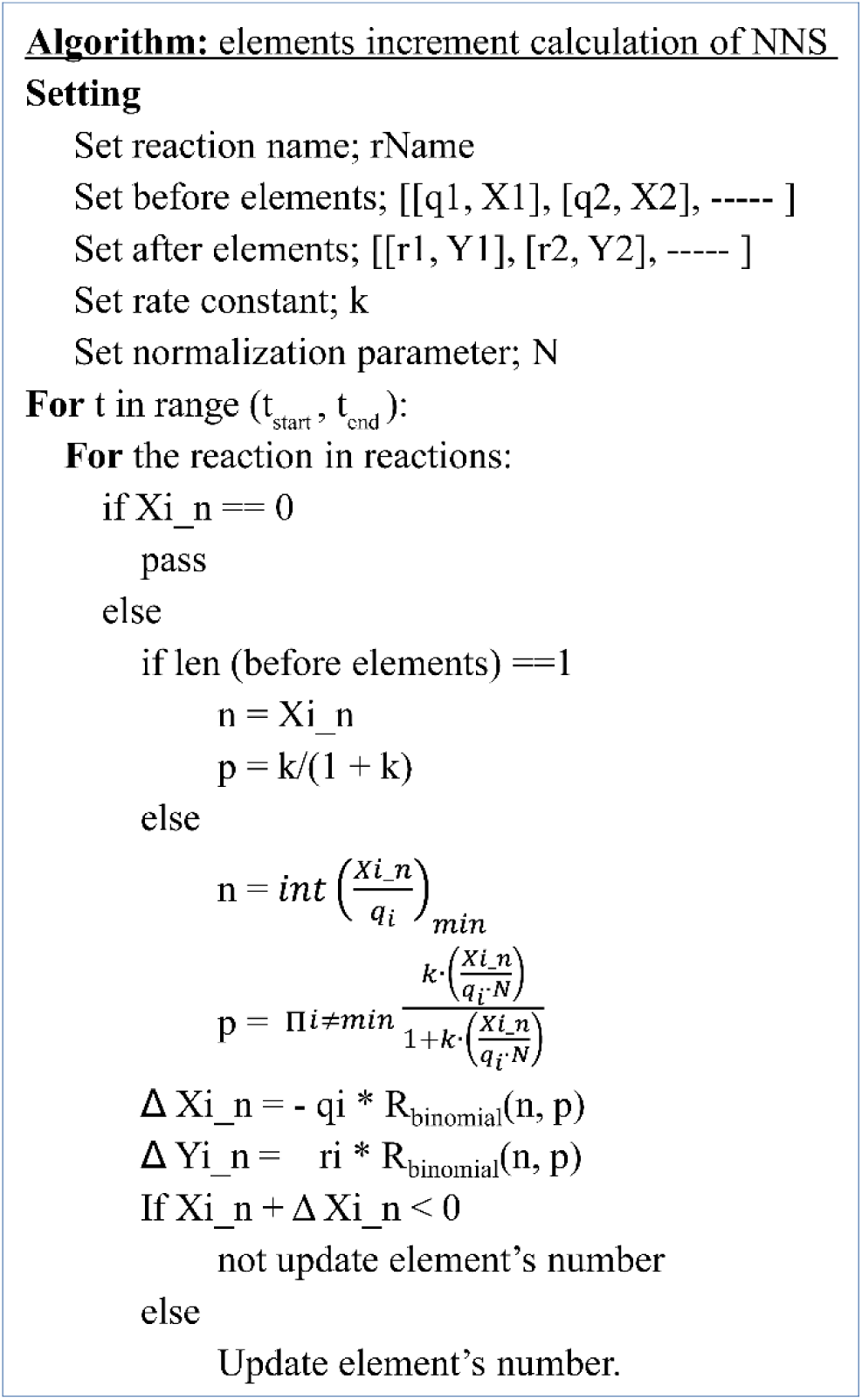
Elements increment algorithm. Main setting and loop.

**Figure S2.**
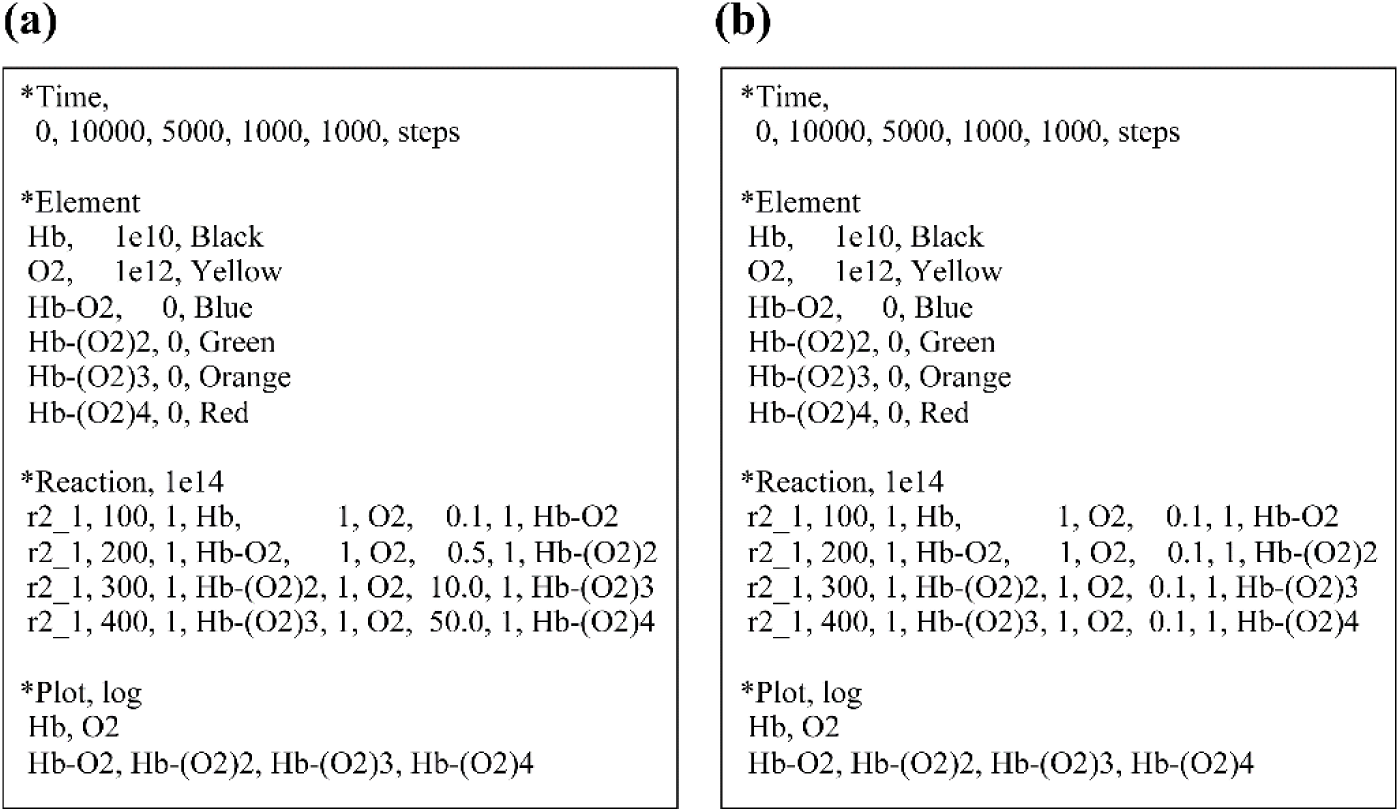
Input text of the Monod–Wyman–Changeux allosteric model. (a) Rate constants increase with O2 binding. (b) Rate constants are equal.

**Figure S3.**
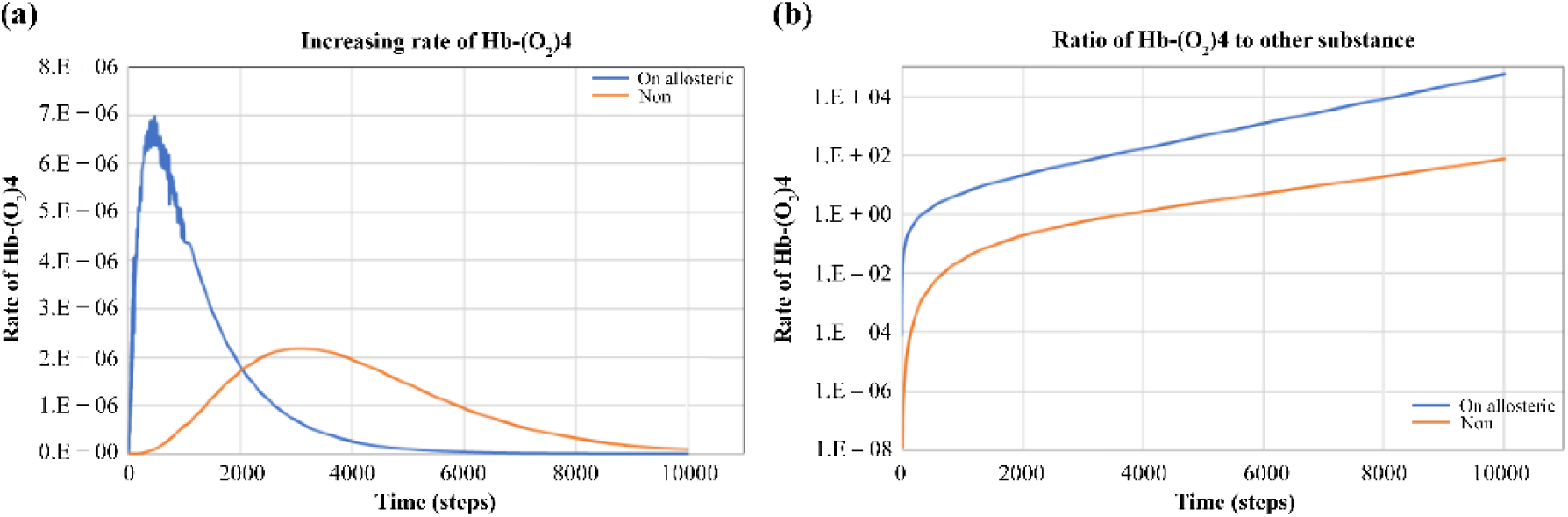
Results of Monod–Wyman–Changeux allosteric model. (a) Increasing rate of Hb-(O_2_)4 of Figure 5. (b) Ratio of Hb-(O2)4 to other substance of Figure 5.

**Figure S4.**
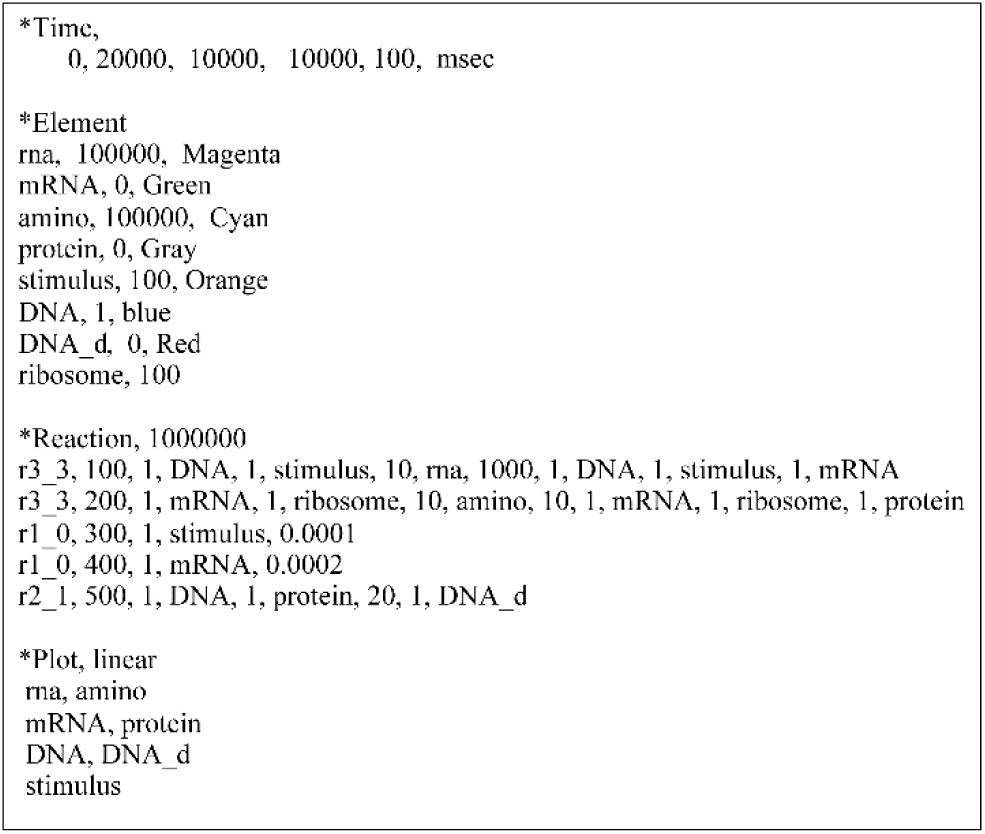
Text input file for the feedback loop.

**Figure S5.**
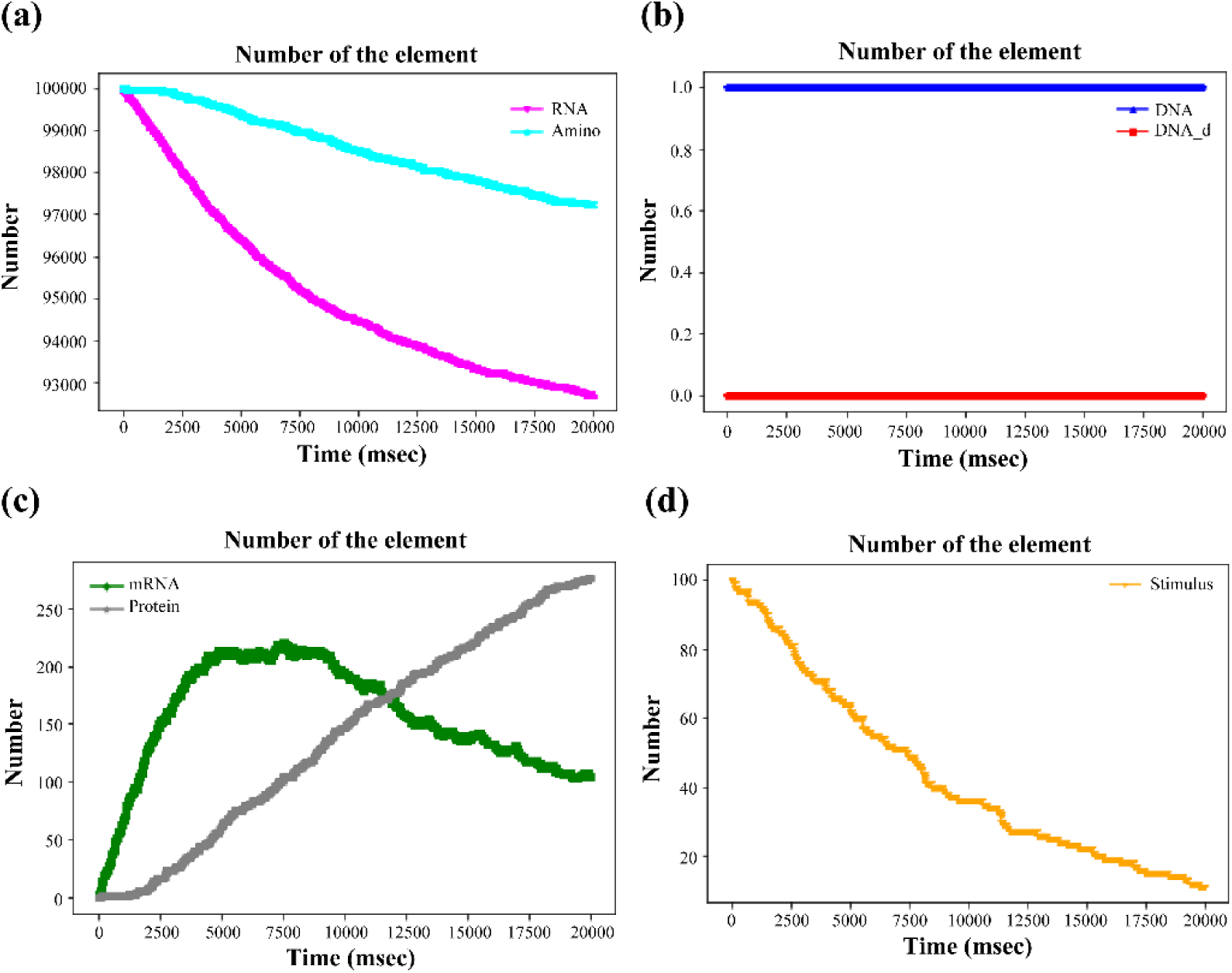
Results of the feedback system of the input file in Supplementary Figure S4 without the reaction “r2_1_500.” (a) RNA and amino acids. (b) DNA and DNA_d. (c) mRNA and protein. (d) Stimulus.

**Figure S6.**
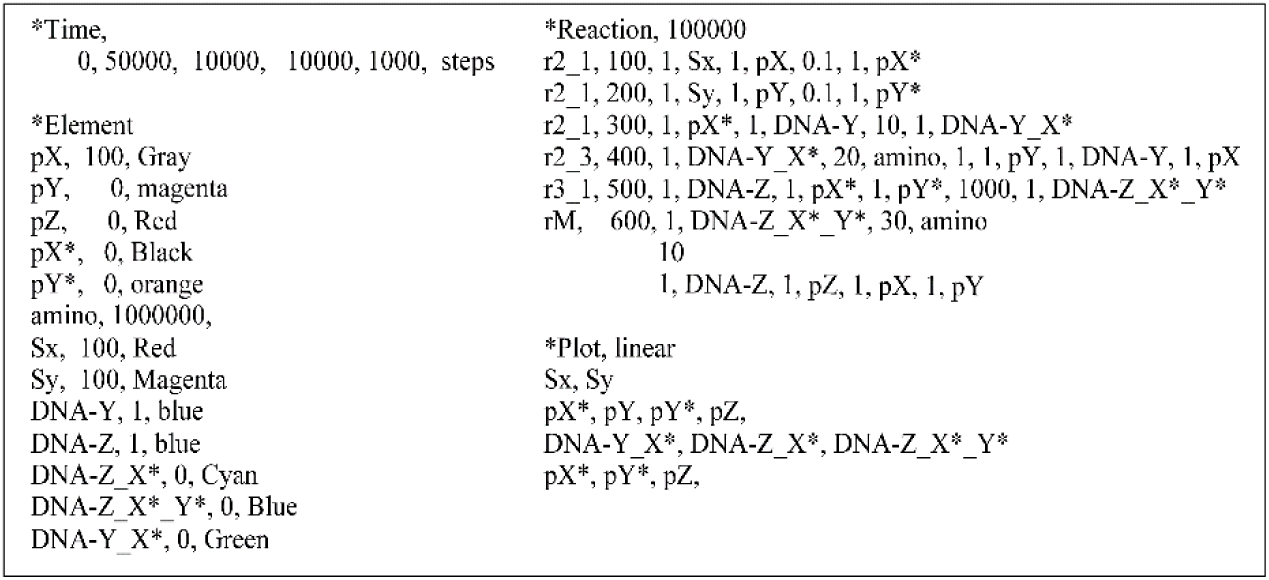
Input file of the feed-forward system in Figure 8.

**Figure S7.**
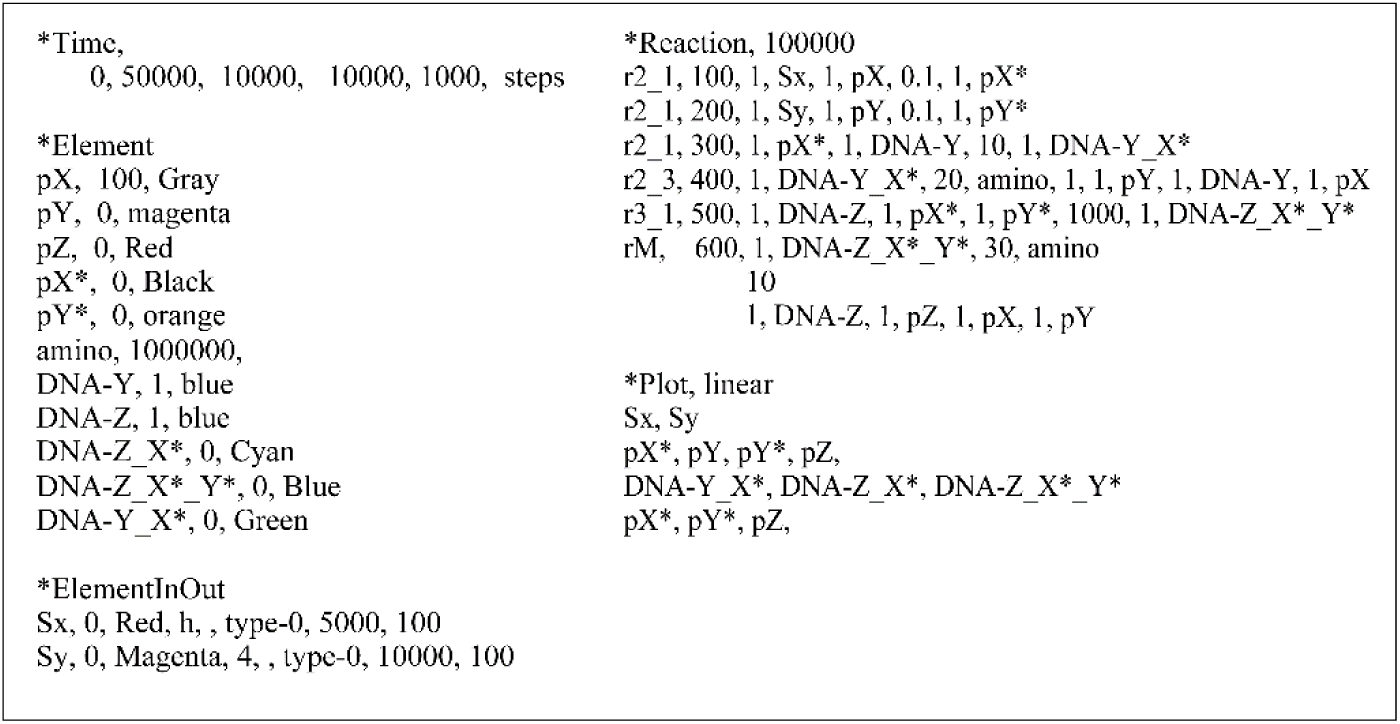
Input file of the feed-forward system. The *Element statements are changed and *ElementInOut in Supplementary Table S3 are added.

**Figure S8.**
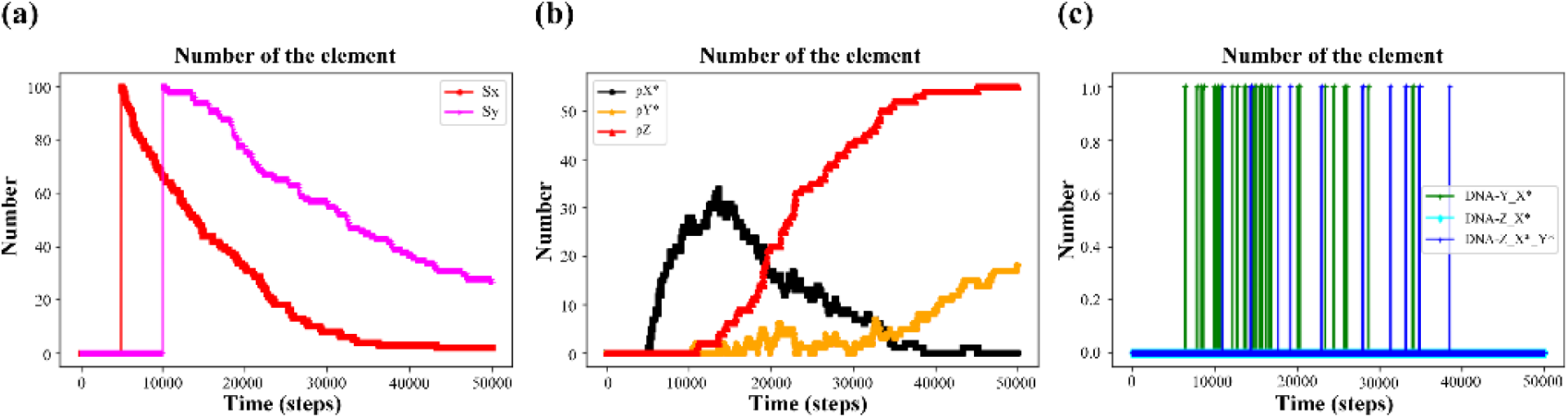
Feed-forward system of Supplementary Figure S7. until time = 50000. Sx and Sy are added at 5000 and 10000, respectively. (a) Sx and Sy. (b) DNA-Y_X*, DNA-Z_X*, and DNA-Z_X*_Y*. (c) pX*, pY*, and pZ.

**Figure S9.**
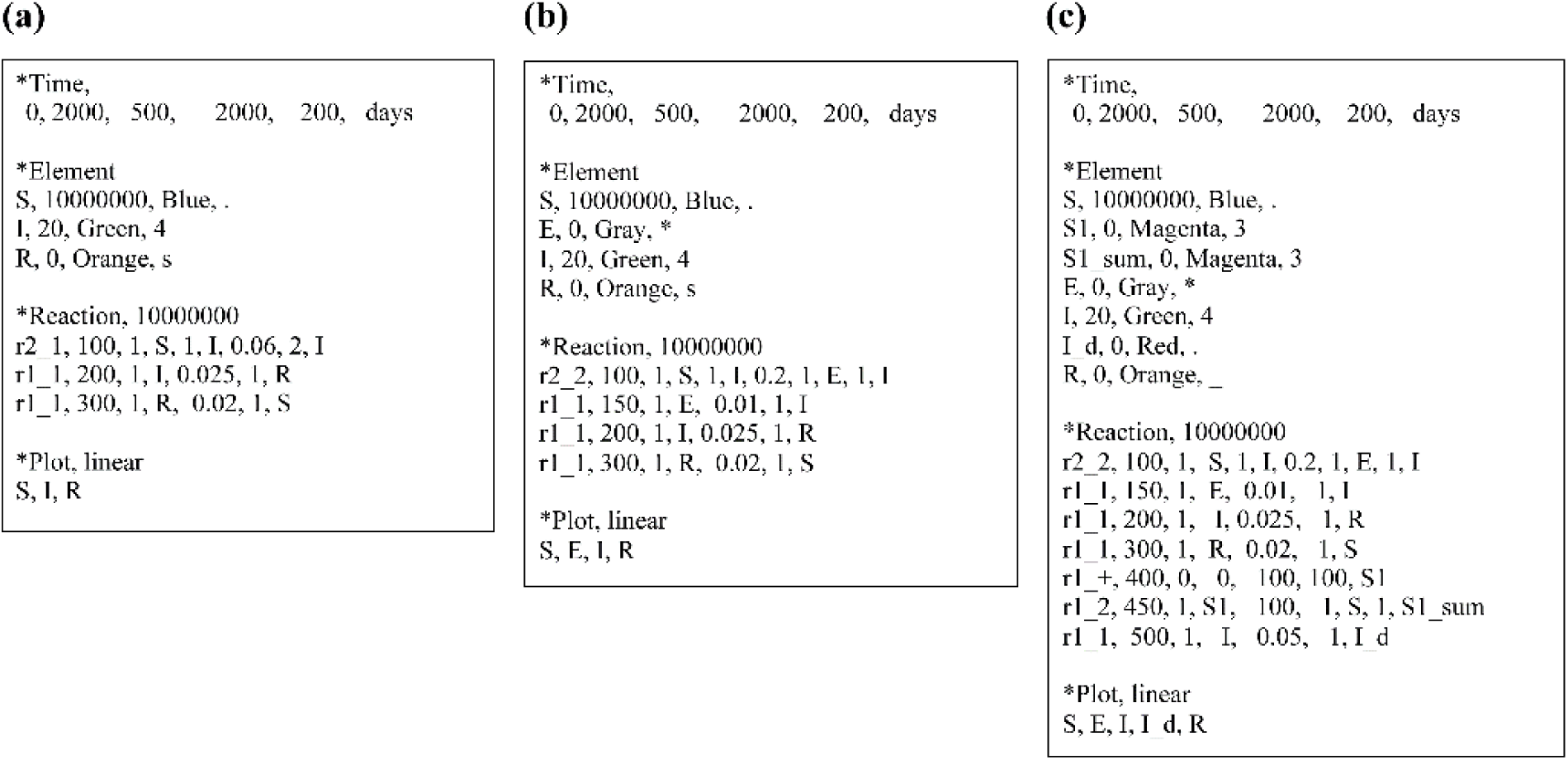
Three input files. (a) SIRS. (b) SEIRS. (c) SEIRS with death and birth.

**Figure S10.**
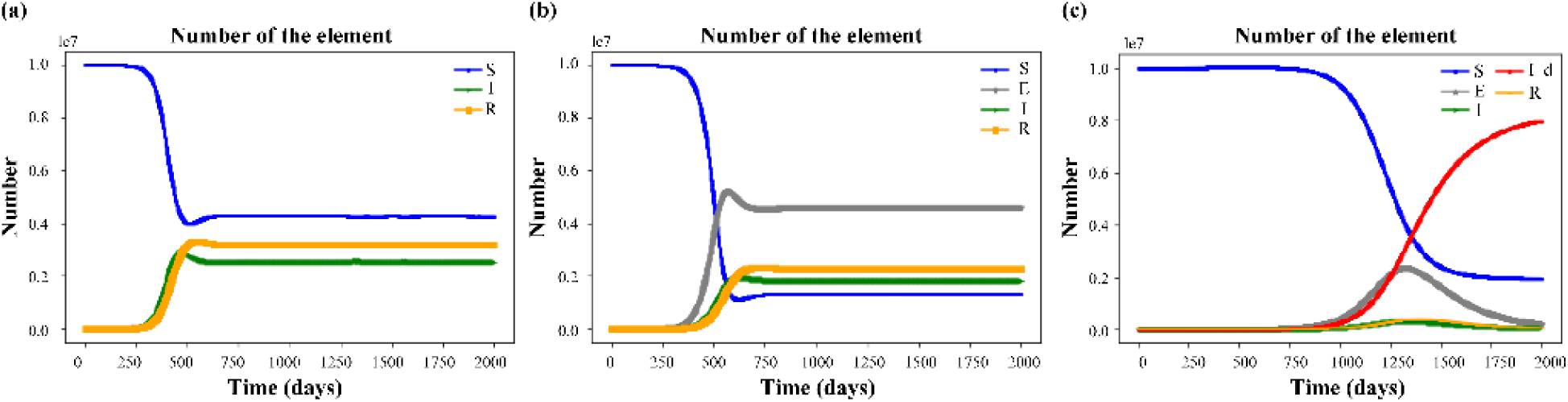
Three results of Supplementary Figure S9. (a) SIRS. (b) SEIRS. (c) SEIRS with death and birth.

**Figure S11.**
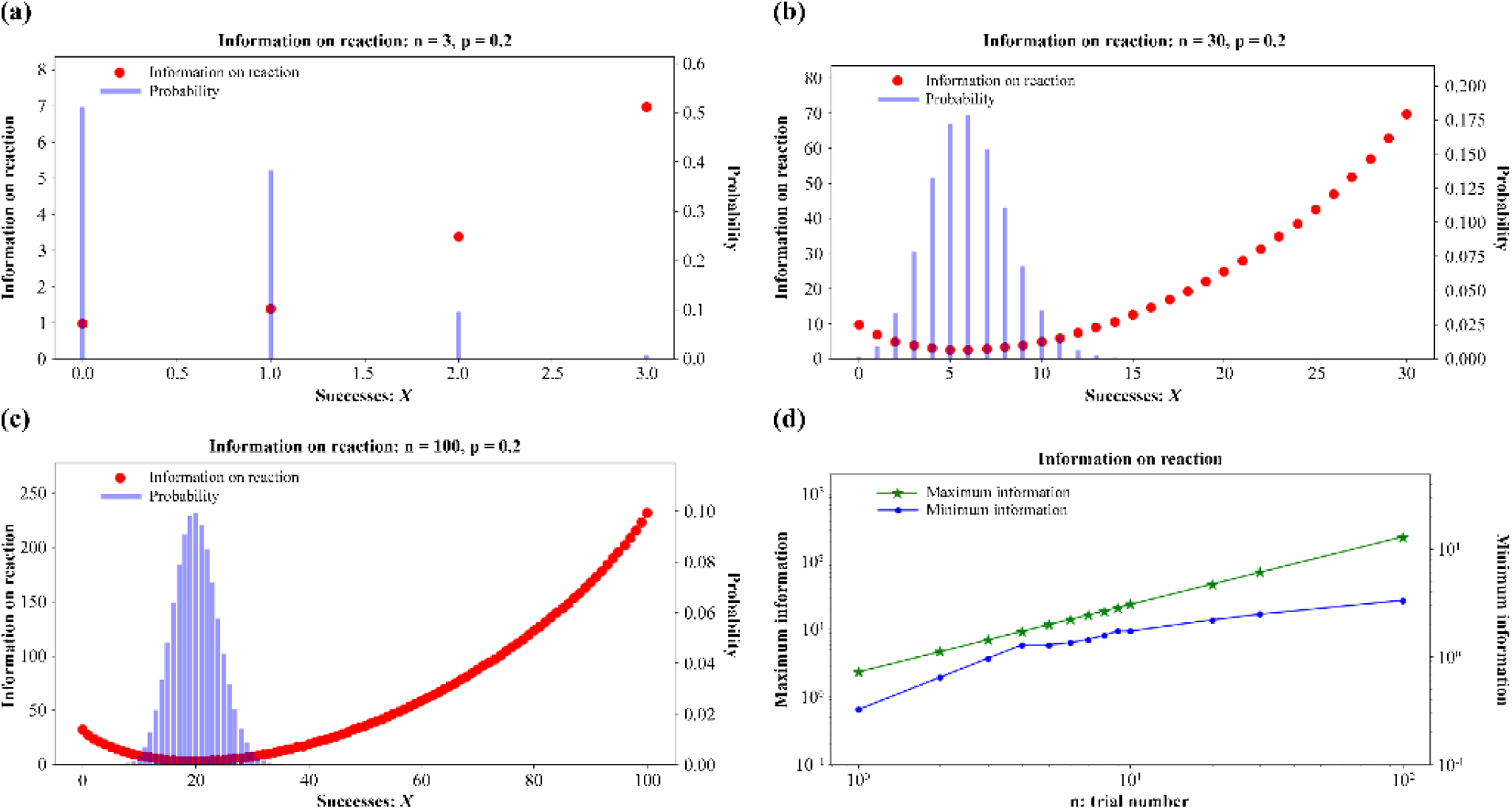
Information on the reaction in Equation (19). (a) n = 3, p = 0.2. (b) n = 30, p = 0.2. (c) n = 300, p = 0.2. (d) Maximum and minimum information on the reaction with p = 0.2.

**Figure S12.**
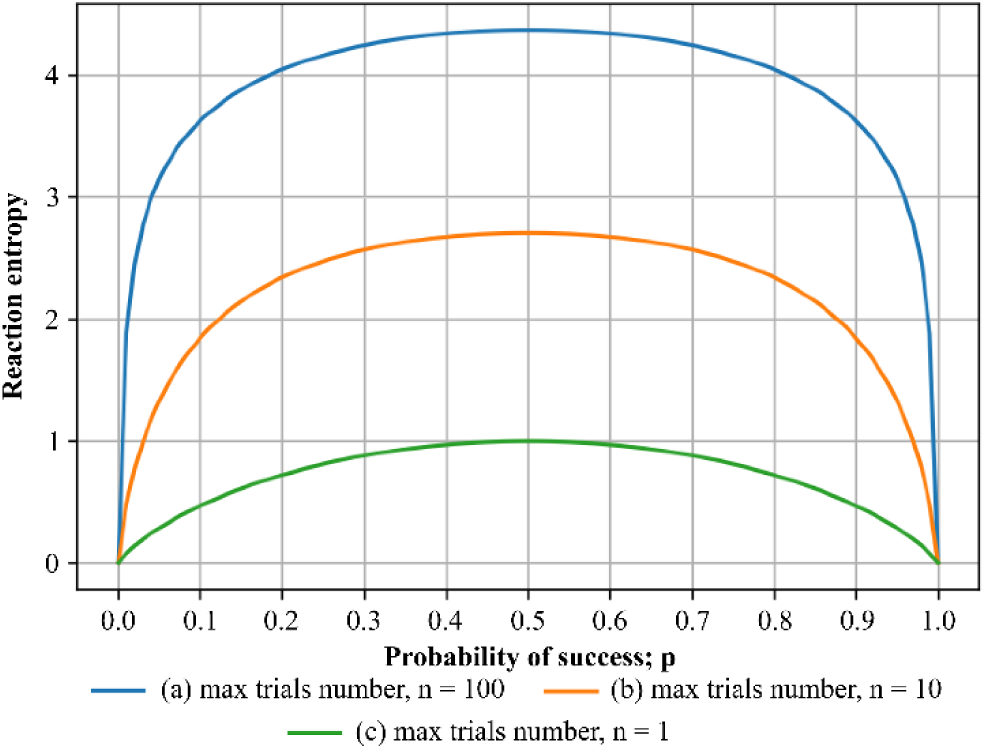
Reaction entropy as a function of p for every three n values.

**Figure S13.**
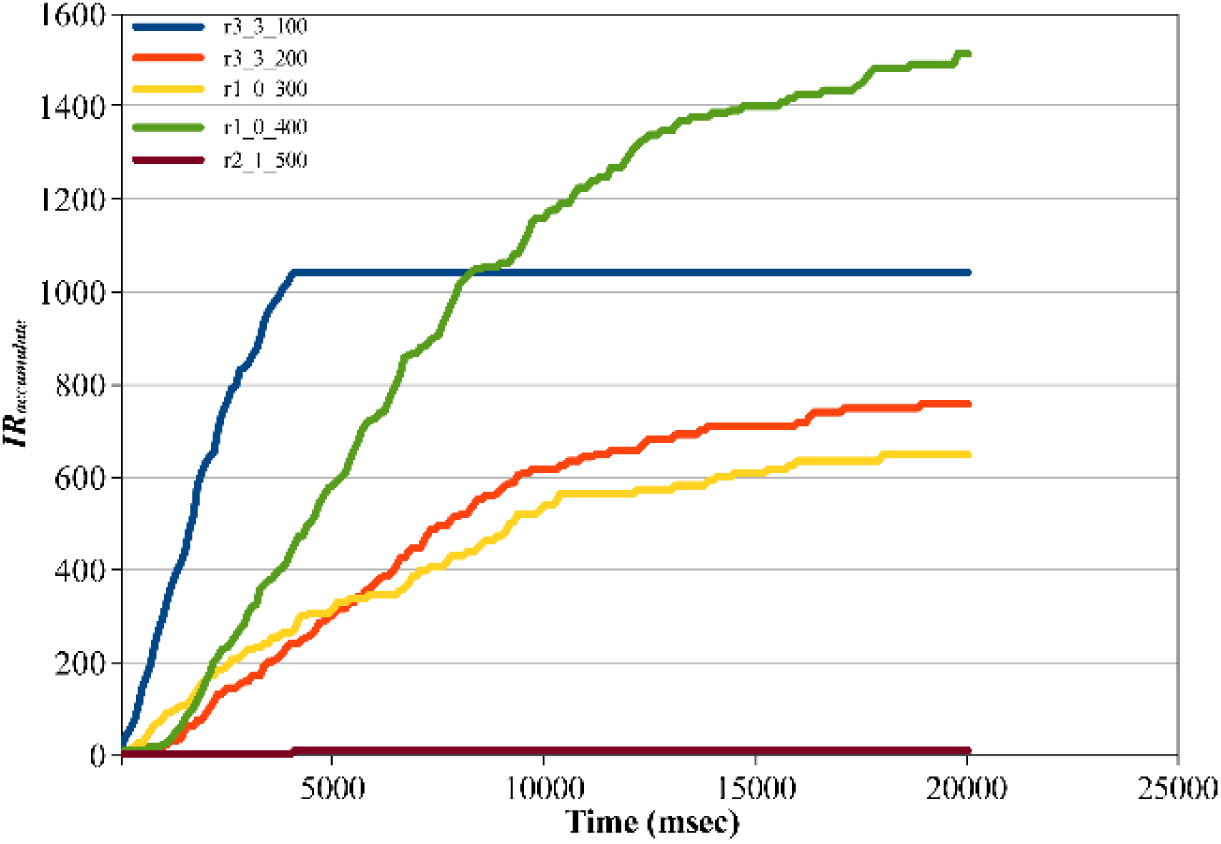
Accumulated information on reaction obtained using Equation (19) in case of a feedback loop of Supplementary Figure S4 with five reactions.

**Figure S14.**
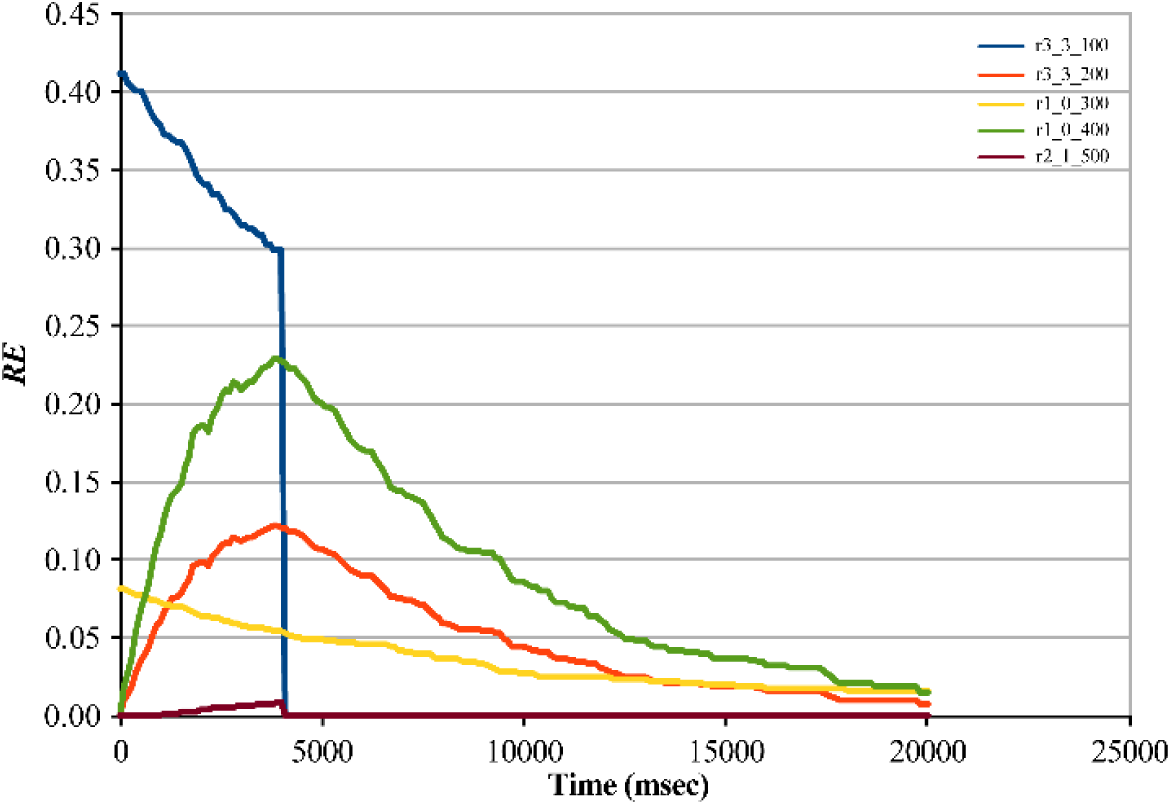
Reaction entropies of Equation (18) at any time in the case of a feedback loop of Supplementary Figure S4 with five reactions.

**Figure S15.**
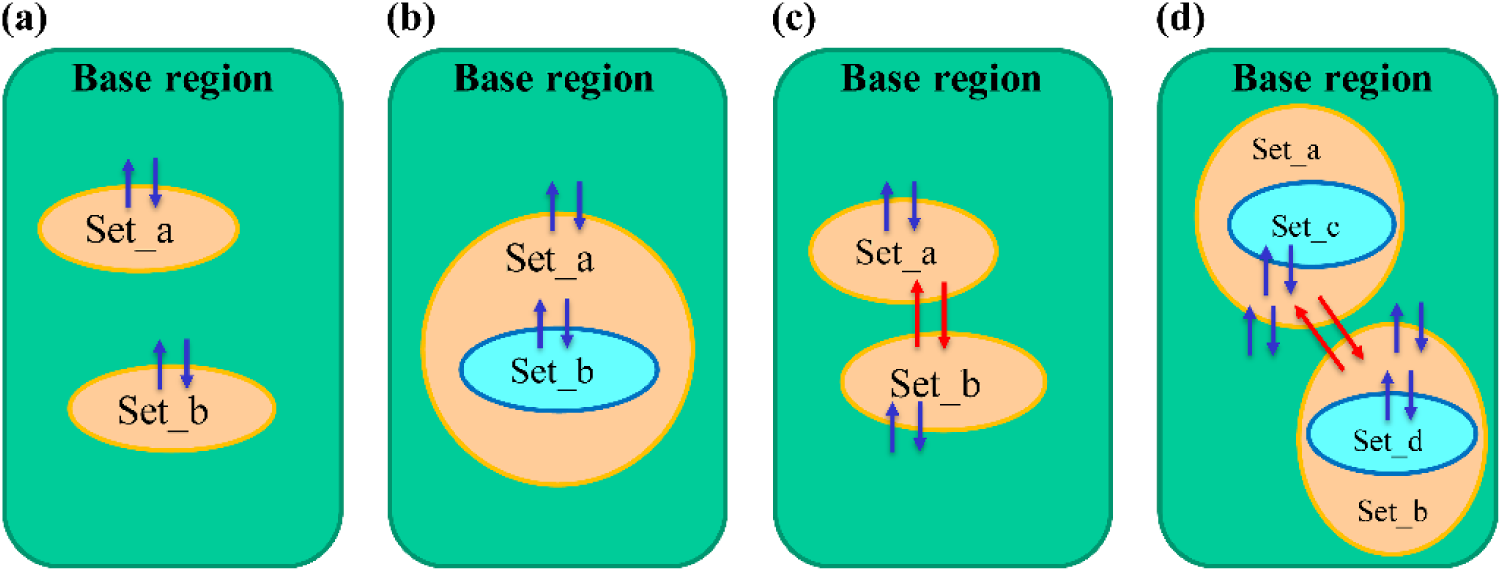
Four types of using “set.” The set contains elements and reactions. The base region also contains elements and reactions. Blue arrows represent translation of elements, and red allows represent direct translation between two sets. (a) Single sets inside a base region. (b) Multilayer modeling. All sets interact with outer region. (c) Direct interaction model between two sets. (d) Direct interaction model between two sets with an inside set each.

